# Comparative analysis of droplet-based ultra-high-throughput single-cell RNA-seq systems

**DOI:** 10.1101/313130

**Authors:** Xiannian Zhang, Tianqi Li, Feng Liu, Yaqi Chen, Jiacheng Yao, Zeyao Li, Yanyi Huang, Jianbin Wang

## Abstract

Since its establishment in 2009, single-cell RNA-seq has been a major driver behind progress in biomedical research. In developmental biology and stem cell studies, the ability to profile single cells confers particular benefits. While most studies still focus on individual tissues or organs, the recent development of ultra-high-throughput single-cell RNA-seq has demonstrated potential power in characterizing more complex systems or even the entire body. However, although multiple ultra-high-throughput single-cell RNA-seq systems have attracted attention, no systematic comparison of these systems has been performed. Here, we focus on three widely used droplet-based ultra-high-throughput single-cell RNA-seq systems, inDrop, Drop-seq, and 10X Genomics Chromium. While each system is capable of profiling single-cell transcriptomes, their detailed comparison revealed the distinguishing features and suitable applications for each system.

## Introduction

Single-cell RNA-seq (scRNA-seq), which was first established in 2009 (Tang et al., 2009), has become one of the most powerful approaches for revealing biological heterogeneity. The ability to manipulate picograms of RNA in single cells has enabled the performance of studies with unprecedented temporal and spatial resolution. Based on the substantial data of the whole transcriptome, scRNA-seq has provided comprehensive information on landscapes of gene expression and their regulatory interactions at the finest resolution, enabling accurate and precise depiction of cell types and states (Grun and van Oudenaarden, 2015; Tanay and Regev, 2017; Wu et al., 2017). In the last decade, the sensitivity and precision of mRNA quantification through scRNA-seq have been greatly improved (Hashimshony et al., 2016; Picelli et al., 2014), leading to revolutionary discoveries in many fields, such as cell-type identification in various tissues or organs (Jaitin et al., 2014; Lake et al., 2016; Papalexi and Satija, 2018; Treutlein et al., 2014; Villani et al., 2017); tracing cell lineage and fate commitment in embryonic development and cell differentiation (Olsson et al., 2016; Semrau et al., 2017; Tirosh et al., 2016; Yan et al., 2013); drawing inferences on transcriptional dynamics and regulatory networks (Deng et al., 2014; Dixit et al., 2016); and identifying the development, evolution, and heterogeneity of tumors (Patel et al., 2014; Treutlein et al., 2014; Venteicher et al., 2017).

The experimental throughput is always a major concern in the design of scRNA-seq experiments. In some biological systems, such as early-stage embryos, only dozens of cells are required to achieve critical findings (Yan et al., 2013). However, owing to tissue complexity, the dynamicity of the cell cycle, or other intrinsic variations (Buettner et al., 2015), as well as technical noise (Brennecke et al., 2013), RNA-seq data from a small number of cells are typically inadequate to reflect the state of biological samples comprehensively (Tanay and Regev, 2017). The sensitivity of transcriptome detection is known to become rapidly saturated with increasing sequencing depth (Svensson et al., 2017). The shallow sequencing of massively sampled single cells can effectively reduce random variation and define cell types through clustering analysis, providing a more robust approach (Pollen et al., 2014; Streets and Huang, 2014; Svensson et al., 2018). For large-scale scRNA-seq studies, a major technical hurdle is the cost of preparing a large number of cDNA libraries. Laboratory automation can overcome the laboriousness of this approach, but the reagents are still expensive (Jaitin et al., 2014). A few recently reported microfluidic approaches have demonstrated various advantages in scRNA-seq (Prakadan et al., 2017). For example, small-volume reactors may improve reaction efficiency and reduce technical noise when coupled with appropriate chemistry (Streets et al., 2014; Wu et al., 2014). Moreover, lab-on-a-chip operations have also made single-cell isolation much easier than manual cell picking (Shalek et al., 2014). Microwell-based scRNA-seq methods (Fan et al., 2015; Han et al., 2018) have also exhibited advantages in terms of low cost and high throughput. However, owing to the lack of commercially available instruments or detailed protocols, microwell-based scRNA-seq has not been widely adopted.

Droplet microfluidics can achieve rapid compartmentation and encapsulation at a frequency of up to dozens of thousands of droplets per second and be easily scaled to produce millions of droplets, each having a nanoliter volume to accommodate single-cell reactions (Agresti et al., 2010). The microfluidic pipeline layout is very simple, consisting mainly of microchannels introducing/collecting reagents and samples (Duncombe et al., 2015). This droplet strategy greatly increases the reaction throughput and dramatically reduces the cost. Currently, there are three prevalent droplet-based systems for high-throughput scRNA-seq, namely, inDrop (Briggs et al., 2018; Klein et al., 2015; Wagner et al., 2018; Zilionis et al., 2017), Drop-seq (Farrell et al., 2018; Macosko et al., 2015), and 10X Genomics Chromium (10X) (Zheng et al., 2017). All of these have been demonstrated to be robust and practical in generating cDNA libraries for thousands of cells in a single run at acceptable cost. All three methods use similar designs to generate droplets, use on-bead primers with barcodes to differentiate individual cells, and apply a unique molecular identifier (UMI) for bias correction (Kivioja et al., 2011). Despite these similarities, they involve different approaches for bead manufacturing, barcode design, and cDNA amplification, and thus have different experimental protocols. Given these differences in system specifications and potentially in the results of transcriptome analysis (Ziegenhain et al., 2017), there is a need for a systematic and unbiased comparison among these methods.

Here, we compare the performance of these three approaches using the same sample with a unified data processing pipeline. We generated two to three replicates for each method using the lymphoblastoid cell line GM12891. The mean sequencing depth was around 50,000–60,000 reads per cell barcode. We also developed a versatile and rapid data processing workflow and applied it for all datasets. Cell capture efficiency, effective read proportion, barcode detection error, and transcript detection sensitivity were analyzed and compared. The results reveal strengths and weaknesses in each system and provide guidance for the selection of the most appropriate system in future research.

## Results

### System overview

Among the three systems, inDrop and Drop-seq have been extensively described in the literature, whereas 10X is a commercial platform whose design details have not been fully disclosed. We here attempt to dissect these systems to the best of our ability based on the information that we could collect. In all three systems, the cell barcodes are embedded in microbead-tethered primers (Figure 1A). The DNA sequences of on-bead primers share a common structure, containing a PCR handle, cell barcode, UMI, and poly-T. The primer on the inDrop beads also has a photo-cleavable moiety and a T7 promoter. However, the beads are fabricated with different materials. The beads used in 10X and inDrop systems are made of hydrogel, while Drop-seq uses brittle resin. Normally, beads and cells are introduced at low concentration to reduce the chance of forming doublets; that is, two cells or two beads are encapsulated in a single droplet. Therefore, for Drop-seq that uses small hard beads, encapsulation of one bead and one cell in the same droplet follows a double Poisson distribution. The hydrogel beads are soft and deformable, closely packed in the microfluidic channel, and their encapsulation can be synchronized to achieve a super-Poissonian distribution (Figure 1A) (Abate et al., 2009). Although 100% single-bead occupancy is very difficult due to inevitable variation in bead size, the cell capture efficiency can reach markedly higher levels in 10X and inDrop approaches. 10X is reported to have ∼80% bead occupancy and a cell capture rate of ∼50% (Zheng et al., 2017).

**Figure 1:**
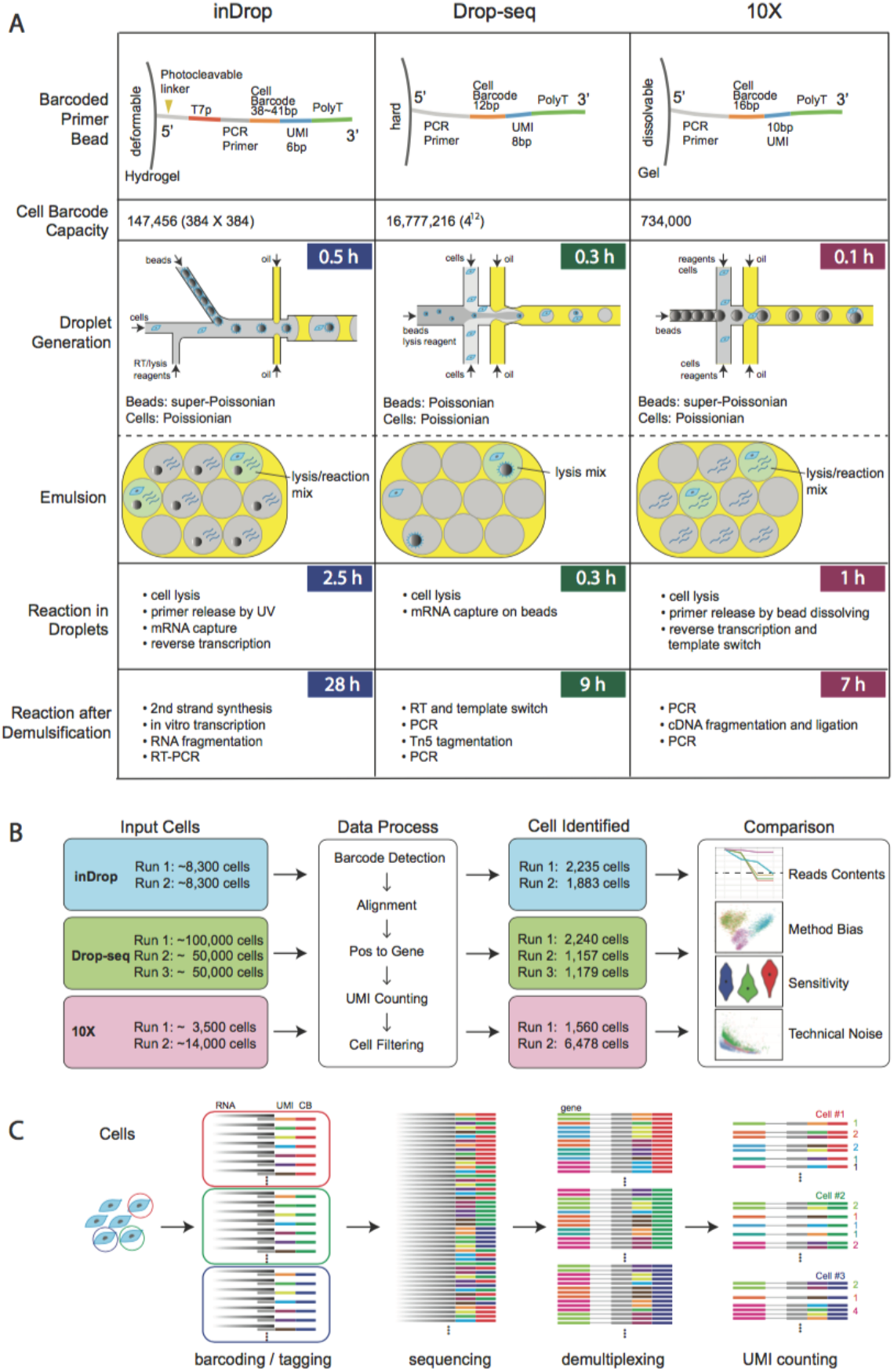
Overview of the three platforms, experimental design, and data analysis pipeline. (A) Schematic and comparison of experimental features of the three systems. They differ in terms of barcode design, library size, emulsion, and downstream reactions. (B) Experimental scheme summary. Two or three replicates were performed for each platform and the same data processing pipeline was used for downstream analysis. The numbers of input and recovered cells are labeled. (C) Overview of the data processing pipeline workflow. The sequencing reads that result from barcoding and tagging in reverse transcription are first demultiplexed by their cell barcodes and then the UMIs mapped to each gene are aggregated and counted.

The material of the beads may also influence the quantity and density of DNA primers. The use of a hydrogel for 10X and inDrop allows the immobilization of primers throughout the beads, whereas the smaller Drop-seq beads can only carry primers on the surface. After encapsulation, the entire beads from 10X are dissolved to release all of the primers into the solution phase to boost the efficiency of mRNA capture. inDrop also mobilizes the primers by UV-irradiation-induced cleavage. In contrast, Drop-seq uses surface-tethered primers to capture the mRNA molecules, which could reduce the capture efficiency compared with that for 10X and inDrop.

Reverse transcription is carried out within droplets for 10X and inDrop before demulsification. Instead, Drop-seq only captures the transcripts without cDNA conversion. Reverse transcription in droplets can confer more uniform results due to the isolation of many local reactions and the reduction of reaction competition. It is also known that the performance of a reaction in a limited volume such as a droplet enhances the specificity of cDNA conversion and relative yield (Streets et al., 2014). The three systems adopt different strategies for cDNA amplification. InDrop employs CEL-Seq (Hashimshony et al., 2012), whereas 10X and Drop-seq follow a template-switching protocol (Macosko et al., 2015; Zheng et al., 2017) similar to the popular Smart-seq chemistry (Ramskold et al., 2012). The *in vitro* transcription step in inDrop extends the library preparation time beyond 24 h, while both Drop-seq and 10X pipelines can be completed within a day.

### Experimental design and data processing

We used GM12891, a human lymphoblastoid cell line, for our comparative study. Biological replicates were set up for all three systems, with various cell inputs on different days and in different batches (Figure 1B). We adjusted the sequencing depth to obtain comparable numbers of reads per cell barcode across the three systems.

Each system has its own data processing pipeline. However, none of them can directly handle data generated by other systems due to differences in read structures. Each analysis pipeline has to cope with system-dependent data characteristics, for example, the tolerance of cell barcode errors. Besides, the analysis pipelines use different strategies in some key processes such as gene tagging. All of these differences may introduce bias in gene quantification, which is not ideal when attempting to perform a fair comparison among the systems. To solve this problem, we developed a versatile pipeline that accepts data from all of these systems and generates matrices of UMI counts (Figure 1C). We applied this pipeline to our data and conducted comparisons on sensitivity, precision, and bias in an objective way.

The script of the pipeline is freely available online (https://github.com/beiseq/baseqDrops) for download. It was designed to accept paired-end sequencing data with one end (read 1) containing the cell barcode and UMI, and the other end (read 2) containing the transcript sequence. The pipeline first identifies cell barcodes in read 1 raw data. After removing cell barcodes with read counts that are too low (miscellaneous barcodes), the pipeline corrects cell barcode errors (see Methods for details). These errors may have been introduced during on-bead primer synthesis and also during PCR or sequencing steps. Reads with the same cell barcodes are aggregated, and invalid cell barcodes are removed after filtering by read counts. For 10X and inDrop in which barcodes are not completely random, the pipeline further filters the cell barcodes based on manufacturers’ whitelists.

Read 2 sequences are mapped to the human reference genome (hg38) using STAR (Dobin et al., 2013) and then tagged to the corresponding genes. We also processed the datasets with each protocol’s official pipeline. We then compared the obtained results with those from our versatile pipeline. The expression levels of the majority of genes and the UMI counts in each barcode were found to be highly consistent among the different data processing methods (see Methods, Figure S2A, B). To confirm the accuracy of transforming aligned reads to the corresponding genes, we performed simulation by generating around 2 million reads based on the cell line’s gene expression profile (ref. f). More than 99% of the reads (2,229,156 out of 2,251,529) were tagged to the correct gene (see Methods, Figure S2C). The remaining 1% of ambiguous reads were mainly derived from genes with paralogs or overlapping genes, such as RPL41/ AC090498.1 or IGHA1/IGHA2 (Table S2). After read-to-gene assignment, the reads for each gene in each cell were grouped and their UMIs were aggregated and counted by allowing a 1-bp mismatch, thus generating a gene expression UMI matrix.

The processing speed of this pipeline was optimized by reducing the read/write payload, which is a common bottleneck. For example, ∼50% of reads from inDrop data have an invalid sequence structure. By removing these reads, we can increase the data processing efficiency. Furthermore, the reads are split into multiple (typically 16) files, based on the cell barcode prefix, which enables parallel processing.

### Quality of primers on beads

The barcode library size determines the maximum capacity for a single experimental run using droplet-based scRNA-seq. A small cell barcode library might result in barcode collision and artificial doublets. In the information accompanying the three systems, theoretical cell barcode library sizes of 1.47×10^5^(inDrop), 1.6×10^7^(Drop-seq), and 7.34×10^5^(10X) are claimed. However, the effective barcode library size may be smaller than the designed value. We estimated the proportion of effective barcodes by analyzing the barcode collisions between multiple runs from each system (see Methods). The likelihood analysis demonstrated the relative probability of observing such a number of collisions at different effective barcode fractions (Figure 2A). For inDrop, our results suggest an effective barcode proportion of around 30%, although 100% effectiveness is also possible with smaller possibility. The analysis is less powerful for larger libraries, but we can still determine the lower bound for Drop-seq (∼10%) and 10X (∼40%). The likelihood of an effective barcode proportion smaller than the lower bound is relatively low. Thus, by rough estimation, the effective barcode size is ∼5×10^4^ for inDrop and at least 1×10^6^ for Drop-seq and 3×10^5^ for 10X (see Methods).

**Figure 2.**
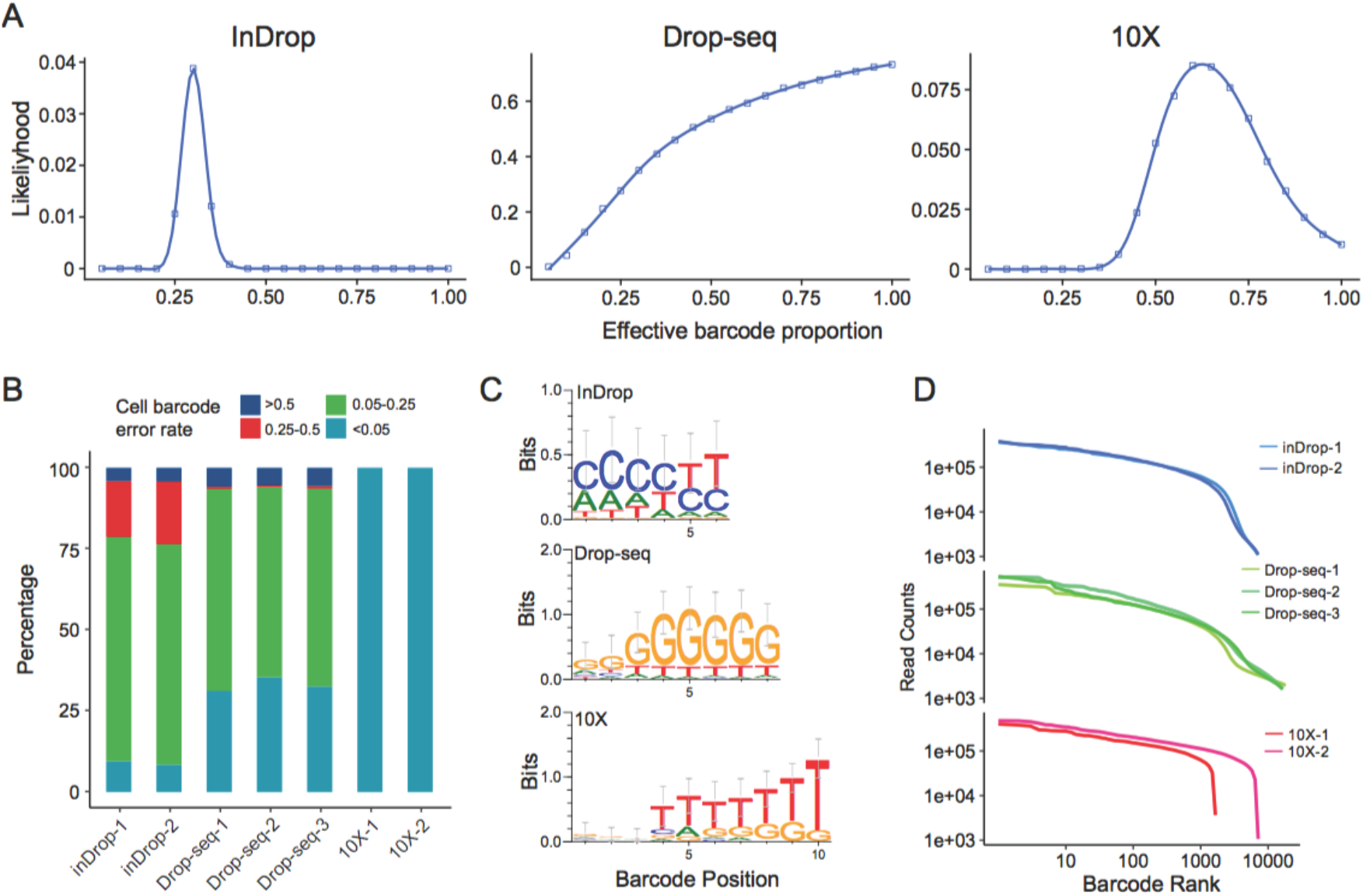
On-bead primer library size and quality assessment. (A) Estimation of effective cell barcode library size for each system. The likelihood of different effective barcode proportion is shown. The likelihood analysis is based on the observed barcode collisions between different samples from the same system (see Methods). (B) Distribution of cell barcode error rate. The error rate was measured as the proportion of corrected reads (1-bp mismatch) relative to the total reads. (C) The motif of the top 50 frequently used UMIs for each system. (D) The primary estimation of the valid cell barcode numbers according to the read counts. Cell barcodes in the same sample are ordered by their read counts. The top N cell barcodes are selected according to input cells and experimental capture efficiency.

One-barcode-one-bead is the key requirement for all three systems. However, owing to the imperfection in the chemistry of DNA synthesis, asynchronous base addition is inevitable. Inconsistency in the sequences of cell barcodes could thus arise within the same bead. Such presence of errors in cell barcodes would result in inflation of the number of detected single cells, which requires careful correction. We aggregated the cell barcodes within 1 Hamming distance. For each valid cell barcode, the proportion of the corrected reads (which contains errors in raw barcode sequences) to the total reads after correction is calculated as the cell barcode error rate (Figure 2B), which reflects the general quality of on-bead DNA primers. 10X beads showed few mismatches in cell barcodes, indicating good quality control in bead fabrication. In contrast, more than half of the cell barcodes contained obvious mismatches in the other two systems. Specifically, about 10% of Drop-seq beads contained a one-base deletion in cell barcodes, which also required extra care during data analysis (see Methods).

We further analyzed the base composition of UMI, which could reflect its synthesis and usage bias (Figure 2C, Table S1). All systems showed bias or preference for poly-T due to its affinity to the poly-A tail of mRNA. We also found the enrichment of poly-C in inDrop, and of poly-G in Drop-seq and 10X. Such patterns, predominantly due to DNA synthesis bias, may cause system-dependent skewness of the RNA-seq results.

The primary filtering criterion for valid cell barcodes is based on the total number of raw reads, which largely reflects the abundance of cellular mRNAs. A cell barcode with more reads is more likely to originate from a real cell. The cell barcodes were sorted and visualized by their read counts, and we observed different features in the three systems (Figure 2D). For 10X, a sharp cliff indicated the distinct difference in read counts between barcodes from healthy cells and others. For inDrop, there was a similar but subtler cliff. For Drop-seq, however, there was no obvious cliff on the read-count curve for a clear cut-off. This might have originated from the wide size distribution of beads used by Drop-seq. We noticed that the size of beads used in inDrop or 10X was more uniform than that in Drop-seq (Figure S1), probably due to the difficulties in size control when fabricating resin beads.

### Data processing steps and results

It is challenging to accurately determine the cell number, represented by cell barcodes, in each sample. This is due to the large dispersion in cellular mRNA molecular counts and their capture efficiency. We attempted multiple strategies to estimate the valid cell numbers (see Methods, Figure S3). Many of these methods rely on certain assumptions about the read/UMI distribution or cell composition, which might not apply for all protocols or situations. We implemented a strategy that started from a certain number of cells determined experimentally, followed by strict quality control filtering (UMIs ≥ 1000 and nearest correlation ≥ 0.6). This strategy has been implemented by multiple groups in recently reported high-throughput scRNA-seq studies. For each run, the number of recovered cells could be roughly estimated by considering the number of input cells, cell capture ratio, and downstream reaction success ratio, in accordance with system-specific protocols. Then, the estimated cells were further filtered to satisfy the quality control criteria (see Methods).

The reads split into each valid cell barcode are first aligned to the human genome to analyze the distribution of reads throughout the genome (Figure 3A). Drop-seq has more than 65% of the reads mapped to UTR (mainly 3’UTR) and exon regions, while this proportion in inDrop is only about 45%. After the tagging of reads that map to gene bodies, the numbers of detectable genes can be obtained (Figure 3B). The number of genes declines in accordance with the number of reads within a cell, except for several outliers in Drop-seq data. We use those detected genes to demonstrate the bias of read distribution along the gene body (Figure 3C). The reads were mainly derived from the 3′ end of the mRNA for all three systems, consistent with their library construction strategies. Drop-seq data showed a bimodal distribution, most likely due to the same PCR anchor sequences being used at both ends of cDNA molecules.

**Figure 3.**
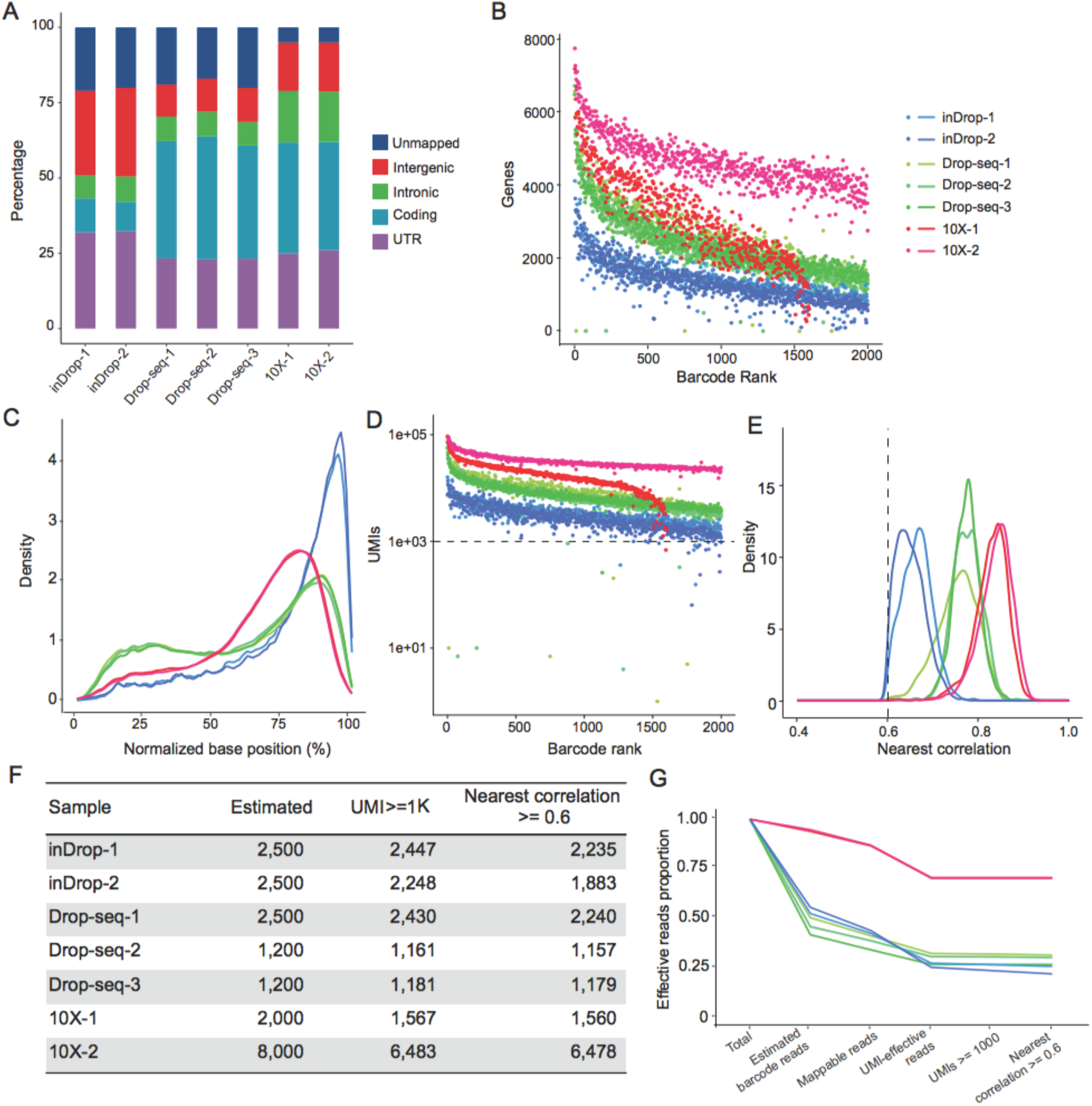
Data processing steps and results. (A) Read composition after mapping to the genome. Percentages of reads mapped to different genomic regions and unmapped reads are shown. (B) The number of genes detected with cell barcode ranked by read counts. (C) Normalized read distribution across the gene body from the 5’ to the 3’ end. (D) The number of UMIs with cell barcode ranked by read counts. (E) The distribution of cells’ nearest correlation (see Methods); a threshold of 0.6 is applied for quality control. (F) The number of valid cell barcodes after each step of quality control filtering. (G) The proportion of effective reads after each step of quality control process (see Methods).

We performed cell barcode filtering based on the total count of UMIs (transcripts) in each experimental run (Figure 3D). With a total UMI cut-off of 1,000, most of the cell barcodes passed the filter, which indicates that the estimated cell number is sound. To further remove possible artifacts caused by barcode errors, we checked the similarity of expression profiles between similar cell barcodes. If the expression profile of a cell barcode was markedly different from its closest cell barcode neighbor (Spearman’s correlation ≤ 0.6, see Methods), we discarded the barcode (Figure 3E, see Methods).

Through all of these steps, we obtained various numbers of cells in each experiment (Figure 3F). The proportion of effective reads (reads from valid barcodes) was ∼75% for 10X, ∼25% for inDrop, and ∼30% for Drop-seq (Figure 3G). The proportion of such reads should be maximized to reduce wastage of sequencing capacity.

### Sensitivity of UMI and gene detection

The sensitivity of gene detection is a fundamental indicator of the performance of scRNA-seq. It reflects the overall efficiency of a method for capturing a single mRNA molecule for reverse transcription, second-strand synthesis, and pre-amplification. It further influences and determines the precision and accuracy of gene expression quantification. With the same cell line as an input sample, the sensitivity can be depicted simply with the recovered UMIs and gene counts (Figure 4A). The UMI and gene numbers gradually become saturated for cell barcodes with increasing read counts (Figure S4A, B). We found that the log-transformed UMI count is highly correlated (Spearman’s correlation r>0.9) with the number of detected genes (Figure S4C). This shows that sequencing depth may influence the numbers of UMIs and genes detected. For a fair comparison among the three different systems, we normalized the dataset to achieve a uniform raw read level (36K/cell) before gene expression analysis (see Methods). The technical replicates from the same system showed high consistency and reproducibility. 10X had the highest sensitivity, capturing over 17,000 transcripts from ∼3,000 genes on average. This performance was consistent regardless of the number of input cells. Drop-seq detected ∼8,000 transcripts from ∼2,500 genes. Meanwhile, the inDrop system had lower sensitivity, detecting ∼2,700 UMIs from ∼1,250 genes. The read distribution is more skewed in inDrop and Drop-seq, for which the majority of cell barcodes have relatively low read counts (Figure 4B).

**Figure 4.**
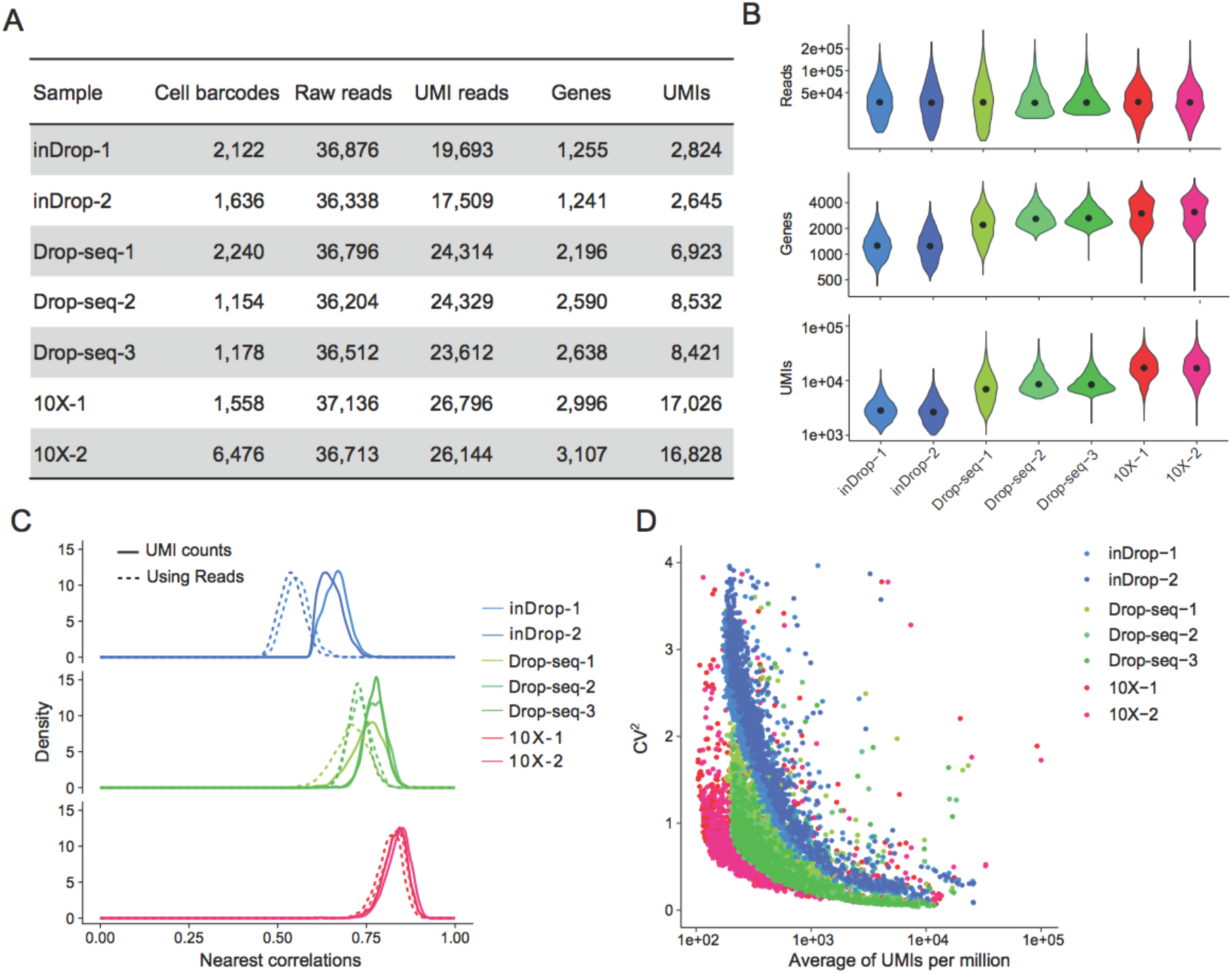
Demonstration of the sensitivity and technical noise of each platform. (A) Summary of cell barcode numbers, read counts, and molecular detection performance. The data are down-sampled to obtain a uniform level of raw reads across all samples (see Methods). (B) The distribution of raw reads, UMIs, and genes detected. (C) Technical noise measured by the nearest correlation between one cell barcode and every other cell barcode within the same sample. Gene quantifications through UMI counts (solid line) and read counts (dashed line) are both adopted. (D) The CV-mean (CV squared) plot of each system. The technical noise is measured at the gene level.

### Technical noise and precision

Technical noise is a reflection of the variation conferred by experimental randomness, including transcript dropout in reverse transcription and the bias associated with PCR amplification. Precision can be assessed by the concordance of the transcriptome among technical replicates. The main purpose of performing single-cell RNA-seq is to cluster cells into different subgroups based on their gene expression profiles, typically for discovering and characterizing new cell types or states. Clustering is based on the similarities or distances of gene expression patterns among cells. Large technical noise or variation will distort the actual distances and obscure subtle biological differences between cells, thus lowering the resolution of cell grouping. Many efforts have been made to reduce the technical noise, such as the use of UMI to eliminate the quantification error caused by amplification bias.

Although we here use an apparently homogeneous cell line, there is still intrinsic biological noise or heterogeneity (Prakadan et al., 2017). In our dataset, the total variation consists of technical and biological components, which are difficult to separate. Here, we assume that biological noise is consistent among samples and that technical noise dominates the variation in the datasets. The noise levels of housekeeping genes (which show a minimal level of biological noise) and other genes have similar distributions, which indicates the low level of biological noise compared with technical noise (Figure S5, see Methods). Thus, the overall total variation should reflect the technical noise level.

The overall total variation is characterized as the nearest Spearman’s correlation between a specific cell barcode and every other cell barcode in the entire dataset (see Methods). Many clustering/classification strategies, such as k-means and hierarchical clustering, are based on the nearest correlation between the cells. To identify minor cell types, the nearest correlation among these minor cells should be high to enable their separation from other cells. To validate the effect of UMIs in reducing the PCR amplification noise of gene counting, we performed the analysis using UMI counts and raw read counts for the quantification of gene expression. The results (Figure 4C) show that 10X and Drop-seq have lower technical noise levels than inDrop. For all three systems, gene expression profiles characterized by UMI have reduced noise compared with those using raw counts, confirming the effectiveness of UMI in noise reduction. It is noteworthy that such noise is more severe in inDrop data, probably due to the use of random primers during library construction. For 10X, however, the usage of UMI does not dramatically reduce noise. This is probably due to relatively even amplification during 10X sample preparation. In addition, most UMIs were sequenced only two to three times, suggesting a less saturated sequencing depth. For deeper sequencing, the use of UMI can probably reduce the noise further.

The technical variation at the gene level can be measured by the coefficient of variation (CV) of normalized UMI (UMIs per million) counts across all cells (Figure 4D, see Methods). This provides a view of the technical noise on the whole gene expression profile. All systems show reduced variation for genes with higher expression levels. Generally, 10X has the lowest technical noise, followed by Drop-seq and then inDrop. Interestingly, many of the most highly expressed genes are quite noisy, especially in the 10X data. We examined these genes (normalized UMI ≥ 2,000, CV ≥ 0.5) and found that most of them were the cell line’s most highly expressed genes or mitochondrial genes (Table S3). High noise in these genes was probably introduced by the stochastic manner of bursts by which transcription occurs (Sanchez and Golding, 2013).

### Saturation of sensitivity and precision at low sequencing depth

The ability to detect transcripts present at a low level could be enhanced by performing deeper sequencing. However, there is a trade-off between costs and sensitivity, especially for high-throughput experiments. Empirically, it has been shown that each cell gets 10,000– 100,000 reads in high-throughput scRNA-seq experiments, whereas for conventional scRNA-seq data the corresponding value is usually ∼1 million reads per cell (Baran-Gale et al., 2017). A previous study based on a mathematical model suggested that shallow sequencing (1% of conventional depth) can also be informative regarding cell status (Heimberg et al., 2016). We randomly subsampled sequencing data and analyzed the corresponding changes in sensitivity and precision (Figures 5A, B and S6). The fitted saturation curves of UMI and gene counts help to determine a suitable sequencing depth for most applications.

**Figure 5.**
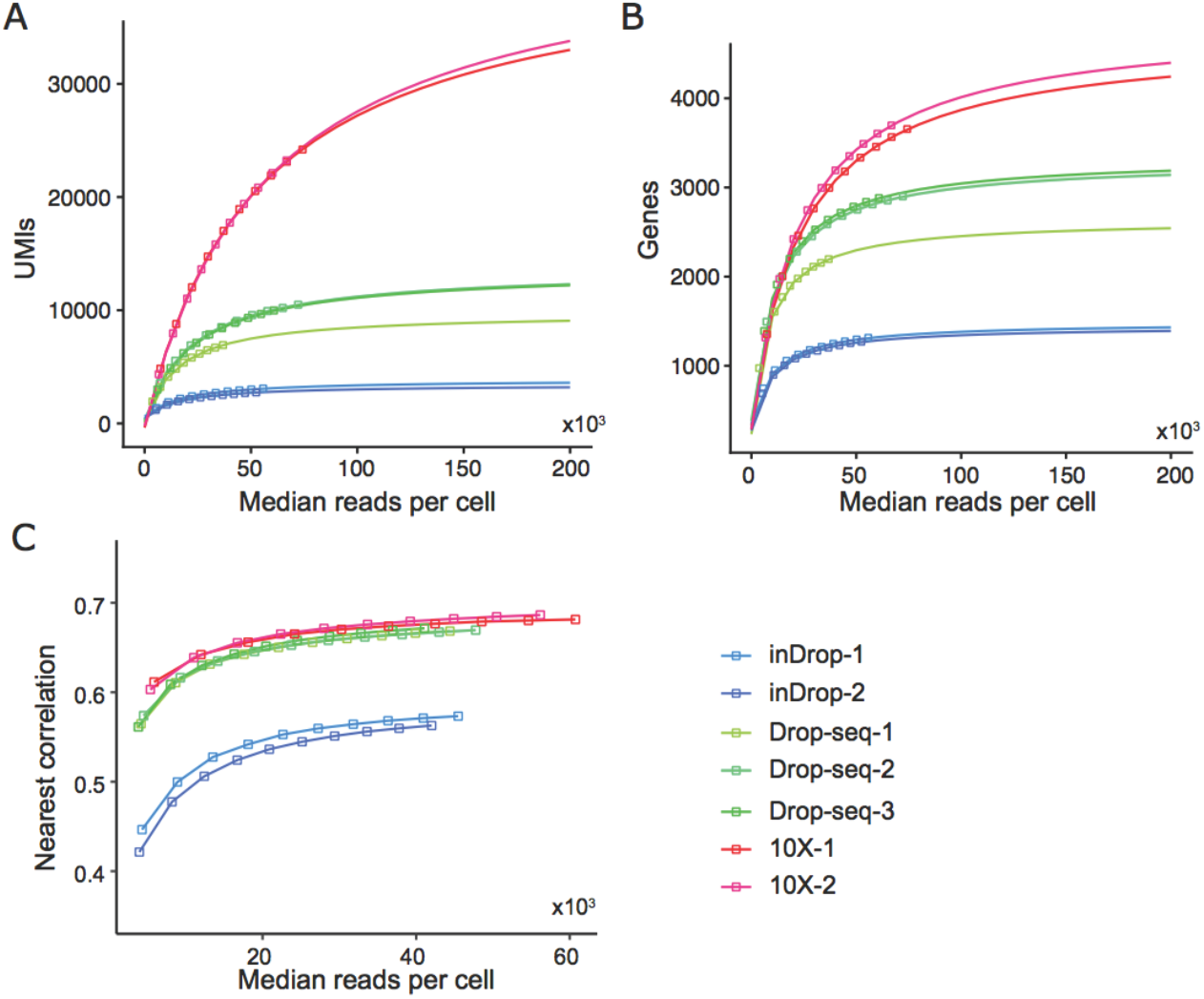
Transcriptome analysis sensitivity and noise level at different sequencing depths by subsampling analysis. Median numbers of UMIs (A) and genes (B) detected for each sample with increasing effective read counts. (C) Transcriptome analysis noise level saturates quickly with sequencing depth. The noise was measured as the nearest correlation (see Methods).

All of the systems show diminishing returns at higher depths. For more sensitive methods, it is possible to detect the same level of UMIs with fewer reads. All three methods can reach a threshold of 1000 UMIs with fewer than 10K reads. 10X can detect 10,000 UMIs with about 20K reads as a median, while for Drop-Seq the value is 50K. These both exceed the capacity of inDrop. We also evaluated how many reads per cell are needed to reach 80% of the total saturated UMIs for Drop-Seq (∼80K) and inDrop (∼60K) (Figure S6A). In contrast, 10X requires ∼200K reads/cell to accomplish this due to the higher sensitivity. Detection sensitivity of gene numbers saturated faster. To reach the 80% saturation level, ∼30K reads/cell are needed for inDrop or Drop-seq, while ∼80K reads/cell are needed for 10X (Figure S6B).

Other than sensitivity, precision also determines a system’s resolution for making biological discoveries. Here, the precision is measured as the nearest correlation between one cell and the others, which also indicates the level of technical noise. We investigated how the precision level was affected by the sequencing depth and found that the precision index rapidly saturated with increasing read depth (≥ 20,000 effective reads) for all three systems (Figure 5C).

These results help us to establish appropriate empirical guidelines for experimental design. For the most commonly performed tasks such as cell typing, a median number of 20,000 reads/cell should be sufficient. However, it should be noted that these results are from a cell line with abundant mRNA. The desired sequencing depth should be considered based on both the sensitivity of protocols and the input RNA content. For cells with lower transcription activities such as primary cells, a lower level of sequencing depth could be sufficient for each protocol.

### Bias in gene quantification

To comprehensively compare the transcriptomes depicted by different systems, we conducted dimension reduction with principal component analysis (PCA) and *t*-distributed stochastic neighbor embedding (tSNE) analyses (Figure 6A). Almost all of the cells were robustly separated and clustered according to their system of origin. Although there is biological and technical variation within cells from the same run, which results in great diversity in sequencing reads, and in gene and UMI counts, the bias between different systems still exceeds the level of these variations. As the replicates are processed in different batches and days, the batch effect is also obscure. Within the same system, different batches of data show a very homogeneous distribution (Figure S7).

**Figure 6.**
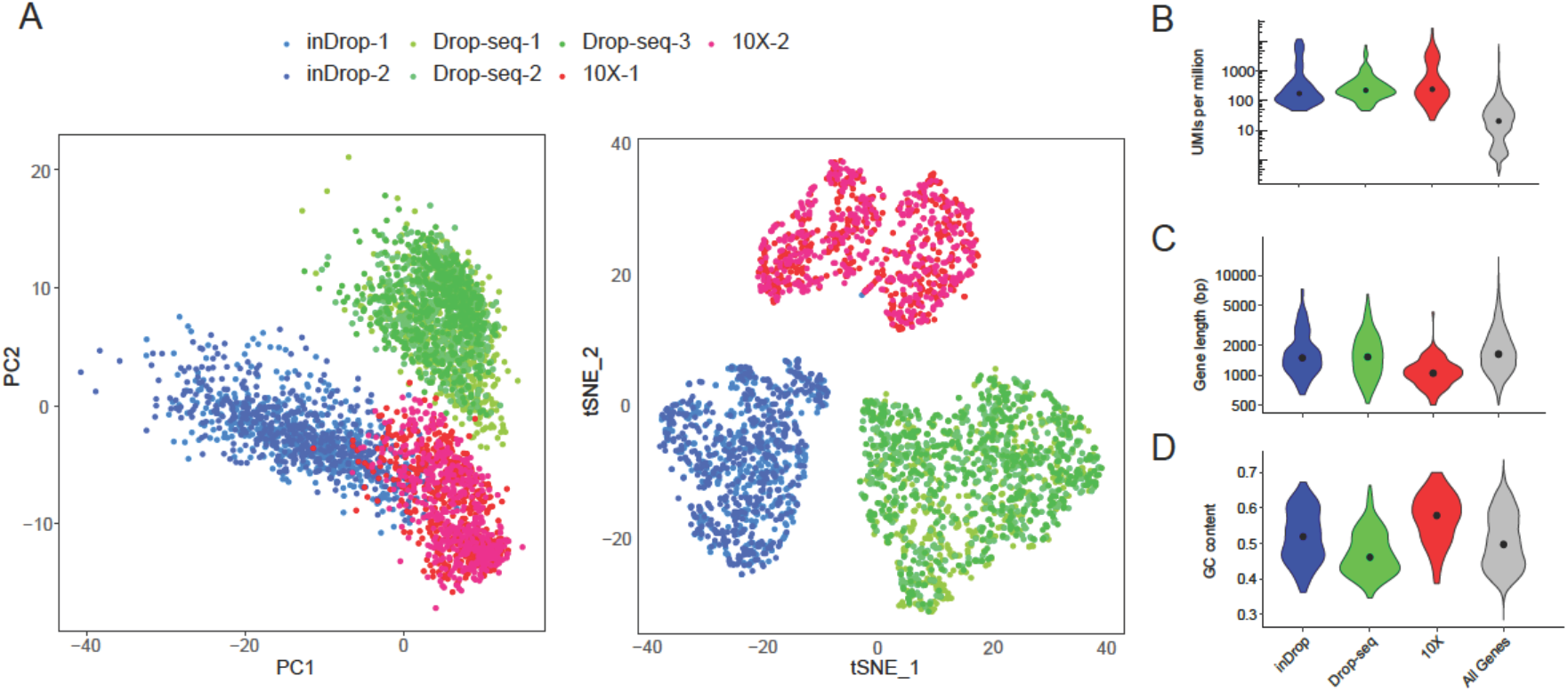
Transcriptome analysis bias in the three systems. (A, B) Visualization of cell barcodes of all three systems clustered by PCA and tSNE. (B–D) Demonstration of transcriptome analysis bias in gene expression level (B), gene length (C), and GC content (D). The top 100 marker genes from each system were used for demonstration.

The separation of cells by system indicates that there is system-specific quantification bias at the gene level. Potential biases in the mRNA enrichment at the gene level could be related to three major factors: expression abundance (normalized to UMIs per million), gene length, and GC content. We hence selected the top 100 marker genes (see Methods) from each method and analyzed the distribution of these factors (Figure 6B–D). These genes showed consistent expression intensity among biological replicates. We found that, compared with the other systems, 10X slightly favored shorter genes and genes with higher GC content, whereas Drop-seq better detected genes with lower GC content. This observation echoes a previous report describing that Drop-seq overestimates transcription of genes with low GC ratio or long sequence (Macosko et al., 2015).

In summary, all of the methods appear to be very consistent and homogeneous among technical replicates from different batches. This indicates the validity of combining different datasets together from the same method. However, different protocols have obvious bias related to gene length and GC content. Thus, combining these datasets directly will introduce extra divergence.

## Discussion

We have compared the three most widely used droplet-based high-throughput single-cell RNA-seq systems, inDrop, Drop-seq, and 10X, using the same cell sample and a unified data processing pipeline to reduce bias in experimental design and data analyses. Technical replicates were included to identify possible batch-dependent artifacts. For each system, we sequenced thousands of single cells. Through quantitative analysis of a few key parameters using our unified data processing pipeline, we have clarified the characteristics of each system. Generally, after filtering out artifacts and errors, all three systems produced quality data for single-cell expression profiling. The cell typing analysis indicated obscure batch effects, but noticeable clustering bias in association with the system of choice. This indicates that cell typing analysis using datasets from a mixture of systems is technically challenging and should be avoided.

In this study, we chose a lymphoblastoid cell line for the analysis because cell line quality is highly controllable. At least for technical evaluation, we wished to reduce the variation of sample quality on the obtained results as much as possible. However, direct comparisons using primary cells, especially those with low mRNA contents, would be more informative. To expand the scope of our study, we further processed HEK293 cells with 10X system and included some datasets produced by the original developers of the three systems (Klein et al., 2015; Macosko et al., 2015; Zheng et al., 2017). As summarized in Table S5, 10X demonstrates higher sensitivity, detecting roughly twice as many of UMIs as inDrop and Drop-seq do from various kinds of cell. The results from the inDrop developers are better than ours. We attribute this discrepancy to batch-to-batch variation in bead quality. As we showed above, inDrop cell barcode error rate is much higher than those of Drop-seq and 10X (Figure 2B). Being labeled with defective barcodes would deem the transcripts undetectable since the very beginning. More than half of inDrop sequencing data were wasted due to a failure of matching with the cell barcodes in our data. Feedback from other inDrop users showed that the equivalent proportions from different batches of beads range from 25% to 65% (unpublished results). We also tested the impact of mRNA content on system performance. When using half of HEK293 cDNA for downstream library preparation, we detected roughly half UMI as in normal HEK293 (Table S5). All these abovementioned results suggest that our findings based on the lymphoblastoid cell line can be generalized to other cell types.

For all three systems, the beads are specifically provided by the particular manufacturer and would probably be difficult to produce in small laboratories. Thus, the quality of the beads, such as their size dispersity, is particularly important to define the robustness and uniformity of reverse transcription and further reactions. Moreover, the fidelity and purity of the barcode sequences on each bead are also key factors affecting the bioinformatics pipeline, for which artifacts and errors should be minimized.

Our comparison shows that 10X generally has higher molecular sensitivity and precision, and less technical noise. As a more maturely commercialized system, the 10X protocol should have been extensively optimized, which is partially reflected in the barcode design and quality control of bead manufacture. However, high-performance optimization also comes with a high price tag. Specifically, the instrument costs more than $50,000, and the per cell cost is around $0.50 even without considering the sequencing cost or instrument depreciation (Table S6).

With small compromises in sensitivity and precision, Drop-seq exhibits a significant advantage in experimental cost compared with 10X, which is typically the major concern when a large number of single cells are needed. As an open-source system (except for the beads), Drop-seq has gained popularity since its introduction in 2015. As of the time of writing, the Drop-seq protocol has been downloaded nearly 60,000 times. Building up the whole system costs less than $30,000. The experimental cost of Drop-seq is about $0.10 per cell (Table S6). Drop-seq is thus a reasonable choice for individual labs given its balanced performance and economical nature.

To a certain extent, inDrop can be considered an open-source version of 10X. Both of them use hydrogel beads for super-Poissonian loading. Their on-bead primers are both releasable to facilitate the capture of transcripts. The instrument cost is comparable to that of 10X, and the per cell cost is about half that of 10X (Table S6). We attribute the lower performance of inDrop to its excessive cDNA amplification (Hashimshony et al., 2016), as well as the fact that the protocol has yet to be completely optimized. As an open-source system, inDrop can adopt other chemistries, and be easily modified for different types of RNA-seq protocols. In a preliminary experiment, we tested the implementation of Smart-seq2, the most widely used scRNA-seq protocol, on the inDrop system. The cDNA profile closely resembled conventional Smart-seq2 products (Figure 7A). We further tested different conditions for reverse transcription and cDNA amplification. Similar to the results generated by the official protocol, a significant proportion (∼40%) of reads in the new data could not be assigned to genuine cell barcodes. Our briefly optimized protocol generated results for UMI and gene detection comparable to those with the official protocol (Figure 7B–D, Table S4). Although the sensitivity of transcription detection was still lower than in the other two systems, our preliminary results demonstrated the flexibility of inDrop, and that the system could be desirable for nonstandard approaches or technical development. With all of the system-specific features mentioned above, we proposed guidance to facilitate the choice of a suitable droplet-based scRNA-seq system for ultra-high-throughput single-cell studies. While most projects work with relatively large cell numbers, precious samples such as human embryos require efficient cell capture. A super-Poissonian distribution of cell capture could be essential for such samples. The requirements regarding the experimental cost and efficiency of transcript detection depend on the specific scenario. Generally, all three systems offer satisfactory transcript detection efficiency, and higher efficiency is associated with higher experimental cost. By rule of thumb, 10X is currently a safe choice for most applications. When the sample is abundant, Drop-seq could be more cost-efficient. In contrast, when the detection of low-abundance transcripts is optional, or a custom protocol is desired, inDrop becomes a better choice.

**Figure 7.**
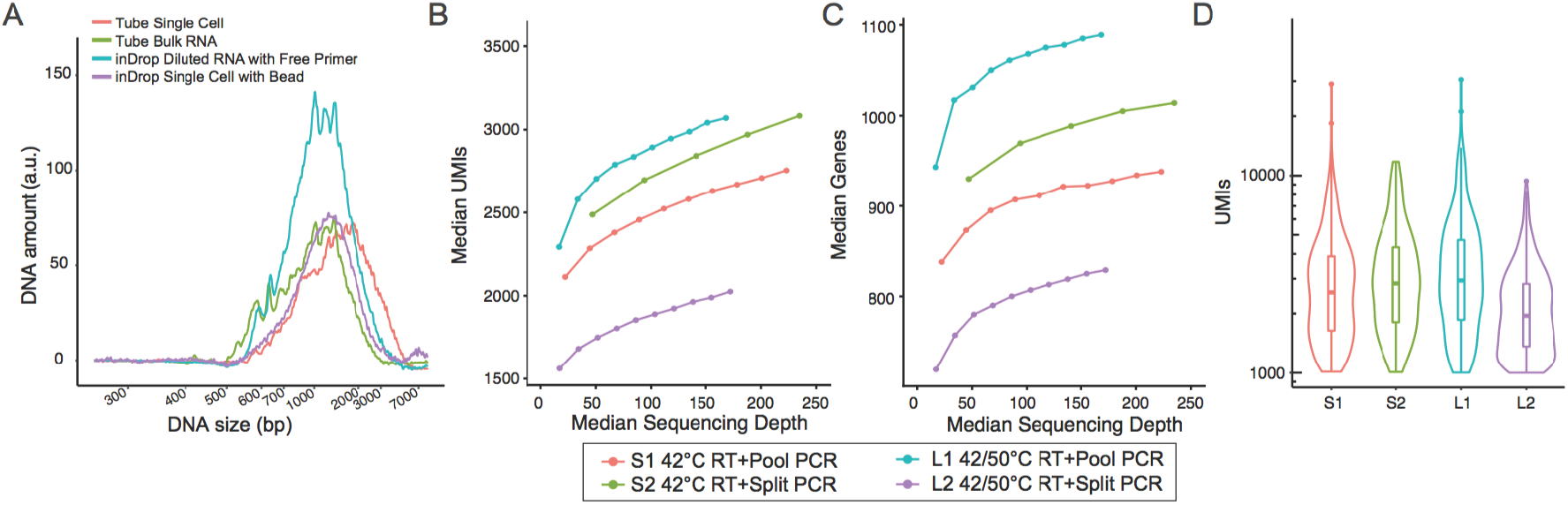
Adopting the Smart-seq2 protocol in the inDrop platform. (A) Comparison of cDNA fragment size between Smart-seq2 performed in tube and inDrop platform. (B, C) Four kinds of reaction with different reaction temperatures and PCR amplification strategies were performed (see Methods). Their median detected UMI (B) and gene (C) counts at various sequencing depths are shown. (D) The UMI distributions for four conditions at uniform sequencing depth (100K reads). The L1 condition has better sensitivity.

## Acknowledgments

We thank Fan Wu for testing the data process pipeline. This work was supported by the National Natural Science Foundation of China (21327808, 21525521 to Yanyi Huang and 21675098 to Jianbin Wang), Ministry of Science and Technology of China (2016YFC0900100 to Jianbin Wang), and Beijing Advance Innovation Center for Genomics.

## Author Contributions

Y.H. and J.W. conceived the project. T.L., F.L., and J.Y. performed experiments. X.Z., Z.L., Y.C., and J.Y. analyzed data. All authors participated in manuscript preparation.

## Data availability

The data can be accessed in GEO using accession code GSE111912.

## References

Abate, A.R., Chen, C.H., Agresti, J.J., and Weitz, D.A. (2009). Beating Poisson encapsulation statistics using close-packed ordering. Lab Chip 9, 2628–2631.

Agresti, J.J., Antipov, E., Abate, A.R., Ahn, K., Rowat, A.C., Baret, J.C., Marquez, M., Klibanov, A.M., Griffiths, A.D., and Weitz, D.A. (2010). Ultrahigh-throughput screening in drop-based microfluidics for directed evolution. Proc Natl Acad Sci U S A 107, 4004–4009.

Baran-Gale, J., Chandra, T., and Kirschner, K. (2017). Experimental design for single-cell RNA sequencing. Brief Funct Genomics.

Brennecke, P., Anders, S., Kim, J.K., Kolodziejczyk, A.A., Zhang, X., Proserpio, V., Baying, B., Benes, V., Teichmann, S.A., Marioni, J.C., et al. (2013). Accounting for technical noise in single-cell RNA-seq experiments. Nat Methods 10, 1093–1095.

Briggs, J.A., Weinreb, C., Wagner, D.E., Megason, S., Peshkin, L., Kirschner, M.W., and Klein, A.M. (2018). The dynamics of gene expression in vertebrate embryogenesis at single-cell resolution. Science.

Buettner, F., Natarajan, K.N., Casale, F.P., Proserpio, V., Scialdone, A., Theis, F.J., Teichmann, S.A., Marioni, J.C., and Stegle, O. (2015). Computational analysis of cell-to-cell heterogeneity in single-cell RNA-sequencing data reveals hidden subpopulations of cells. Nat Biotechnol 33, 155–160.

Deng, Q., Ramskold, D., Reinius, B., and Sandberg, R. (2014). Single-cell RNA-seq reveals dynamic, random monoallelic gene expression in mammalian cells. Science 343, 193–196.

Dixit, A., Parnas, O., Li, B., Chen, J., Fulco, C.P., Jerby-Arnon, L., Marjanovic, N.D., Dionne, D., Burks, T., Raychowdhury, R., et al. (2016). Perturb-Seq: Dissecting Molecular Circuits with Scalable Single-Cell RNA Profiling of Pooled Genetic Screens. Cell 167, 1853– 1866 e1817.

Dobin, A., Davis, C.A., Schlesinger, F., Drenkow, J., Zaleski, C., Jha, S., Batut, P., Chaisson, M., and Gingeras, T.R. (2013). STAR: ultrafast universal RNA-seq aligner. Bioinformatics 29, 15–21.

Duncombe, T.A., Tentori, A.M., and Herr, A.E. (2015). Microfluidics: reframing biological enquiry. Nat Rev Mol Cell Biol 16, 554–567.

Fan, H.C., Fu, G.K., and Fodor, S.P. (2015). Expression profiling. Combinatorial labeling of single cells for gene expression cytometry. Science 347, 1258367.

Farrell, J.A., Wang, Y., Riesenfeld, S.J., Shekhar, K., Regev, A., and Schier, A.F. (2018). Single-cell reconstruction of developmental trajectories during zebrafish embryogenesis. Science.

Grun, D., and van Oudenaarden, A. (2015). Design and Analysis of Single-Cell Sequencing Experiments. Cell 163, 799–810.

Han, X., Wang, R., Zhou, Y., Fei, L., Sun, H., Lai, S., Saadatpour, A., Zhou, Z., Chen, H., Ye, F., et al. (2018). Mapping the Mouse Cell Atlas by Microwell-Seq. Cell 172, 1091–1107 e1017.

Hashimshony, T., Senderovich, N., Avital, G., Klochendler, A., de Leeuw, Y., Anavy, L., Gennert, D., Li, S., Livak, K.J., Rozenblatt-Rosen, O., et al. (2016). CEL-Seq2: sensitive highly-multiplexed single-cell RNA-Seq. Genome Biol 17, 77.

Hashimshony, T., Wagner, F., Sher, N., and Yanai, I. (2012). CEL-Seq: single-cell RNA-Seq by multiplexed linear amplification. Cell Rep 2, 666–673.

Heimberg, G., Bhatnagar, R., El-Samad, H., and Thomson, M. (2016). Low Dimensionality in Gene Expression Data Enables the Accurate Extraction of Transcriptional Programs from Shallow Sequencing. Cell Syst 2, 239–250.

Jaitin, D.A., Kenigsberg, E., Keren-Shaul, H., Elefant, N., Paul, F., Zaretsky, I., Mildner, A., Cohen, N., Jung, S., Tanay, A., et al. (2014). Massively parallel single-cell RNA-seq for marker-free decomposition of tissues into cell types. Science 343, 776–779.

Kivioja, T., Vaharautio, A., Karlsson, K., Bonke, M., Enge, M., Linnarsson, S., and Taipale, J. (2011). Counting absolute numbers of molecules using unique molecular identifiers. Nat Methods 9, 72–74.

Klein, A.M., Mazutis, L., Akartuna, I., Tallapragada, N., Veres, A., Li, V., Peshkin, L., Weitz, D.A., and Kirschner, M.W. (2015). Droplet barcoding for single-cell transcriptomics applied to embryonic stem cells. Cell 161, 1187–1201.

Lake, B.B., Ai, R., Kaeser, G.E., Salathia, N.S., Yung, Y.C., Liu, R., Wildberg, A., Gao, D., Fung, H.L., Chen, S., et al. (2016). Neuronal subtypes and diversity revealed by single-nucleus RNA sequencing of the human brain. Science 352, 1586–1590.

Macosko, E.Z., Basu, A., Satija, R., Nemesh, J., Shekhar, K., Goldman, M., Tirosh, I., Bialas, A.R., Kamitaki, N., Martersteck, E.M., et al. (2015). Highly Parallel Genome-wide Expression Profiling of Individual Cells Using Nanoliter Droplets. Cell 161, 1202–1214.

Olsson, A., Venkatasubramanian, M., Chaudhri, V.K., Aronow, B.J., Salomonis, N., Singh, H., and Grimes, H.L. (2016). Single-cell analysis of mixed-lineage states leading to a binary cell fate choice. Nature 537, 698–702.

Papalexi, E., and Satija, R. (2018). Single-cell RNA sequencing to explore immune cell heterogeneity. Nat Rev Immunol 18, 35–45.

Patel, A.P., Tirosh, I., Trombetta, J.J., Shalek, A.K., Gillespie, S.M., Wakimoto, H., Cahill, D.P., Nahed, B.V., Curry, W.T., Martuza, R.L., et al. (2014). Single-cell RNA-seq highlights intratumoral heterogeneity in primary glioblastoma. Science 344, 1396–1401.

Picelli, S., Faridani, O.R., Bjorklund, A.K., Winberg, G., Sagasser, S., and Sandberg, R. (2014). Full-length RNA-seq from single cells using Smart-seq2. Nat Protoc 9, 171–181.

Pollen, A.A., Nowakowski, T.J., Shuga, J., Wang, X., Leyrat, A.A., Lui, J.H., Li, N., Szpankowski, L., Fowler, B., Chen, P., et al. (2014). Low-coverage single-cell mRNA sequencing reveals cellular heterogeneity and activated signaling pathways in developing cerebral cortex. Nat Biotechnol 32, 1053–1058.

Prakadan, S.M., Shalek, A.K., and Weitz, D.A. (2017). Scaling by shrinking: empowering single-cell ’omics’ with microfluidic devices. Nat Rev Genet 18.

Ramskold, D., Luo, S., Wang, Y.C., Li, R., Deng, Q., Faridani, O.R., Daniels, G.A., Khrebtukova, I., Loring, J.F., Laurent, L.C., et al. (2012). Full-length mRNA-Seq from single-cell levels of RNA and individual circulating tumor cells. Nat Biotechnol 30, 777–782.

Sanchez, A., and Golding, I. (2013). Genetic determinants and cellular constraints in noisy gene expression. Science 342, 1188–1193.

Semrau, S., Goldmann, J.E., Soumillon, M., Mikkelsen, T.S., Jaenisch, R., and van Oudenaarden, A. (2017). Dynamics of lineage commitment revealed by single-cell transcriptomics of differentiating embryonic stem cells. Nat Commun 8, 1096.

Shalek, A.K., Satija, R., Shuga, J., Trombetta, J.J., Gennert, D., Lu, D., Chen, P., Gertner, R.S., Gaublomme, J.T., Yosef, N., et al. (2014). Single-cell RNA-seq reveals dynamic paracrine control of cellular variation. Nature 510, 363–369.

Streets, A.M., and Huang, Y. (2014). How deep is enough in single-cell RNA-seq? Nat Biotechnol 32, 1005–1006.

Streets, A.M., Zhang, X., Cao, C., Pang, Y., Wu, X., Xiong, L., Yang, L., Fu, Y., Zhao, L., Tang, F., et al. (2014). Microfluidic single-cell whole-transcriptome sequencing. Proc Natl Acad Sci U S A 111, 7048–7053.

Svensson, V., Natarajan, K.N., Ly, L.H., Miragaia, R.J., Labalette, C., Macaulay, I.C., Cvejic, A., and Teichmann, S.A. (2017). Power analysis of single-cell RNA-sequencing experiments. Nat Methods 14, 381–387.

Svensson, V., Vento-Tormo, R., and Teichmann, S.A. (2018). Exponential scaling of single-cell RNA-seq in the past decade. Nat Protoc 13, 599–604.

Tanay, A., and Regev, A. (2017). Scaling single-cell genomics from phenomenology to mechanism. Nature 541, 331–338.

Tang, F., Barbacioru, C., Wang, Y., Nordman, E., Lee, C., Xu, N., Wang, X., Bodeau, J., Tuch, B.B., Siddiqui, A., et al. (2009). mRNA-Seq whole-transcriptome analysis of a single cell. Nat Methods 6, 377–382.

Tirosh, I., Venteicher, A.S., Hebert, C., Escalante, L.E., Patel, A.P., Yizhak, K., Fisher, J.M., Rodman, C., Mount, C., Filbin, M.G., et al. (2016). Single-cell RNA-seq supports a developmental hierarchy in human oligodendroglioma. Nature 539, 309–313.

Treutlein, B., Brownfield, D.G., Wu, A.R., Neff, N.F., Mantalas, G.L., Espinoza, F.H., Desai, T.J., Krasnow, M.A., and Quake, S.R. (2014). Reconstructing lineage hierarchies of the distal lung epithelium using single-cell RNA-seq. Nature 509, 371–375.

Venteicher, A.S., Tirosh, I., Hebert, C., Yizhak, K., Neftel, C., Filbin, M.G., Hovestadt, V., Escalante, L.E., Shaw, M.L., Rodman, C., et al. (2017). Decoupling genetics, lineages, and microenvironment in IDH-mutant gliomas by single-cell RNA-seq. Science 355.

Villani, A.C., Satija, R., Reynolds, G., Sarkizova, S., Shekhar, K., Fletcher, J., Griesbeck, M., Butler, A., Zheng, S., Lazo, S., et al. (2017). Single-cell RNA-seq reveals new types of human blood dendritic cells, monocytes, and progenitors. Science 356.

Wagner, D.E., Weinreb, C., Collins, Z.M., Briggs, J.A., Megason, S.G., and Klein, A.M. (2018). Single-cell mapping of gene expression landscapes and lineage in the zebrafish embryo. Science.

Wu, A.R., Neff, N.F., Kalisky, T., Dalerba, P., Treutlein, B., Rothenberg, M.E., Mburu, F.M., Mantalas, G.L., Sim, S., Clarke, M.F., et al. (2014). Quantitative assessment of single-cell RNA-sequencing methods. Nat Methods 11, 41–46.

Wu, A.R., Wang, J., Streets, A.M., and Huang, Y. (2017). Single-Cell Transcriptional Analysis. Annu Rev Anal Chem (Palo Alto Calif) 10, 439–462.

Yan, L., Yang, M., Guo, H., Yang, L., Wu, J., Li, R., Liu, P., Lian, Y., Zheng, X., Yan, J., et al. (2013). Single-cell RNA-Seq profiling of human preimplantation embryos and embryonic stem cells. Nat Struct Mol Biol 20, 1131–1139.

Zheng, G.X., Terry, J.M., Belgrader, P., Ryvkin, P., Bent, Z.W., Wilson, R., Ziraldo, S.B., Wheeler, T.D., McDermott, G.P., Zhu, J., et al. (2017). Massively parallel digital transcriptional profiling of single cells. Nat Commun 8, 14049.

Ziegenhain, C., Vieth, B., Parekh, S., Reinius, B., Guillaumet-Adkins, A., Smets, M., Leonhardt, H., Heyn, H., Hellmann, I., and Enard, W. (2017). Comparative Analysis of Single-Cell RNA Sequencing Methods. Mol Cell 65, 631–643 e634.

Zilionis, R., Nainys, J., Veres, A., Savova, V., Zemmour, D., Klein, A.M., and Mazutis, L. (2017). Single-cell barcoding and sequencing using droplet microfluidics. Nat Protoc 12, 44– 73.

